# Inducible depletion of Scleraxis-lineage cells during tendon healing impairs multi-scale restoration of tendon structure

**DOI:** 10.1101/2022.03.17.484720

**Authors:** Antonion Korcari, Samantha Muscat, Mark R. Buckley, Alayna E. Loiselle

## Abstract

Tendons are composed of a heterogeneous cell environment, with *Scleraxis*-lineage (*Scx*^*Lin*^) cells being the predominant population. Although *Scx*^*Lin*^ cells are required for maintenance of tendon homeostasis, their functions during tendon healing are unknown. To this end, we first characterized the spatiotemporal dynamics of *Scx*^*Lin*^ cells during tendon healing, and identified that the overall *Scx*^*Lin*^ pool continuously expands up to early remodeling healing phase. To better define the function of *Scx*^*Lin*^ cells during the late proliferative phase of healing, we inducibly depleted *Scx*^*Lin*^ cells from day 14-18 post-surgery using the Scx-Cre; Rosa-DTR mouse model, with local administration of diphtheria toxin inducing apoptosis of *Scx*^*Lin*^ cells in the healing tendon. At D28 post-surgery, *Scx*^*Lin*^ cell depleted tendons (DTR) had substantial impairments in structure and function, relative to WT, demonstrating the importance of *Scx*^*Lin*^ cells during tendon healing. Next, bulk RNAseq was utilized to identify the underlying mechanisms that were impaired with depletion and revealed that *Scx*^*Lin*^ depletion induced molecular and morphological stagnation of the healing process at D28. However, this stagnation was transient, such that by D56 tendon mechanics in DTR were not significantly different than wildtype repairs. Collectively, these data offer fundamental knowledge on the dynamics and roles of *Scx*^*Lin*^ cells during tendon healing.

## Introduction

Tendons are dense fibrous tissues that connect muscles to bones to enable skeletal movement and joint stability [1]. Such forces can be transmitted successfully due to tendon’s structure, which consists of a highly aligned and organized Collagen I rich matrix. Acute tendon injuries are significant since they account for approximately 30% of all musculoskeletal consultations and despite great advances in surgical and rehabilitation protocols, 30-40% of flexor tendon repairs still result in unsatisfactory outcomes [2–4]. After injury, tendons heal via the production of fibrotic scar tissue, characterized by abundant and disorganized extracellular matrix (ECM) [5–11]. Despite efforts to regain pre-injury structure via an on-going remodeling process, the imposed structural and functional deficits are largely permanent. Currently, there are signfiicant knowledge gaps related to the specific cell populations and biological mechanisms responsible for tendon healing [12], and as such, there is a paucity of pharmacological strategies to improve the healing process.

While recent studies have demonstrated heterogeneity of the adult tendon cell environment [7, 13, 14], nearly all cells in the adult tendon are encompassed by the Scleraxis (*Scx*)-lineage [7], with *Scx*, a basic helix-loop-helix transcription factor, being the most-well characterized tendon marker [15]. Recent work has demonstrated that the overall Scx-lineage pool (*Scx*^*Lin*^, as labelled using the non-inducible Scx-Cre) can be further broken down into different subpopulations during post-natal growth, adult homeostasis, and phases of the tendon healing process as shown using the Scx-Cre^ERT2^ driver [13], while other studies have demonstrated *de novo* activation of *Scx* in response to injury [16]. However, whether these subpopulations make distinct contributions to the healing process are unknown. To begin to address this, we recently demonstrated that depletion of *Scx*^*Lin*^ cells from adult tendon prior to injury improved the healing process [7]. Intriguingly, lineage tracing suggested that there was a new population of *Scx*^*+*^ cells that was transiently added to the overall *Scx*^*Lin*^ pool by D14 post-surgery such that *Scx*^*Lin*^ depletion prior to injury had minimal effect on the early phases of healing. In contrast, the impact of *Scx*^*Lin*^ depletion prior to injury was more profound during late healing (D28) with a concomitant reduction in the overall *Scx*^*Lin*^pool, and functional improvements in healing. Taken together, although the above studies have shed some light on the potential temporally-distinct roles of the overall *Scx*^*Lin*^ pool in tendon healing, there are still significant knowledge gaps in terms of their temporal dynamics and time-dependent functions during tendon healing.

In this study, our first goal was to comprehensively define the dynamics of *Scx*^*Lin*^ cells during tendon healing. In specific, we focused on both adult *Scx*^*Lin*^ cells (cells that expressed *Scx* during adult tendon homeostasis) as well as cells that express *Scx* in response to the injury using the Scx-GFP reporter. Second, we hypothesized that *Scx*^*Lin*^ cells contribute to the bridging of tendon stubs via ECM synthesis, and thus, *Scx*^*Lin*^ depletion during the proliferative healing phase will significantly impair the composition, structure, and function of the newly formed bridging tissue. Deciphering the time-dependent roles of *Scx*^*Lin*^ cells during tendon healing will better inform therapeutic target selection and promote regenerative tendon healing.

## Materials and methods

### Mice

*Scx-Cre (*MGI:5317938), *Scx-Cre*^*ERT2*^ [8, 17] and *Scx*^*GFP*^ [18, 19] mice were generously provided by Dr. Ronen Schweitzer. Rosa-Ai9 (#007909) and Rosa-DTR^LSL^ (#007900) mice were obtained from the Jackson Laboratory (Bar Harbor, ME, USA). For the lineage-tracing studies, first, *Scx-Cre*^*ERT2*^ mice were crossed to Rosa-Ai9 to generate *Scx-Cre*^*ERT2,Ai9*^ mice. Next, homozygous *Scx-Cre*^*ERT2,Ai9*^ mice were crossed to *Scx*^*GFP/GFP*^ to generate *Scx-Cre*^*ERT2,Ai9*^*;Scx*^*GFP*^ mice. These mice received three 100 mg/kg intraperitoneal (i.p.) tamoxifen (TMX) injections beginning seven days prior to FDL tendon injury and repair surgery to label and trace cells that actively expressed *Scx* during homeostasis (adult *Scx*^*Lin*^ cells), while ensuring that no subsequent labeling occurred during FDL tendon healing by introducing a washout period of four days between the last i.p. TMX injection and the tendon surgeries. In parallel, the GFP reporter in these mice allows for additional labeling and tracing of all cells that actively express *Scx* at the time of harvest. For the depletion studies, *Scx-Cre* mice were crossed to the Rosa-DTR^LSL^ mice to generate *Scx-Cre*^*+*^; *DTR*^*F/+*^, a model of broad *Scx*^*Lin*^ cell depletion (DTR) [7, 13], while *Scx-Cre*^*-*^; *DTR*^*F/+*^ mice were used as wildtype littermates controls (WT). Diphtheria toxin receptor (Rosa-DTR^LSL^) mice can be utilized to temporally ablate cell populations in a cell/tissue-type specific manner using Cre drivers [20]. The expression of DTR is inhibited prior to Cre-mediated recombination due to the presence of a STOP cassette flanked by loxp site (Loxp-STOP-Loxp; LSL). The STOP cassette is deleted due to the Cre-mediated recombination, which results in the expression of the DTR. In this case, DTR is expressed in the overall *Scx*^*Lin*^ pool. Administration of diphtheria toxin (DT) to these mice results in targeted cell death of the broad *Scx*^*Lin*^ pool. All mouse studies were performed with 10–12 week-old male and female mice. All mouse work (injections, surgeries, harvests) were performed in the morning. Mice were kept in a 12 hr light/dark cycle.

### Flexor tendon repair

At 10–12 weeks of age, mice underwent complete transection and repair of the flexor digitorum longus (FDL) tendon in the hind paw as previously described [5]. Briefly, mice received an injection of sustained-release buprenorphine (1mg/kg). Next, they were anesthetized with Ketamine (60 mg/kg) and Xylazine (4 mg/kg). To reduce chances of rupture at the repair site, the FDL tendon was first transected at the myotendinous junction and the skin was closed with a 5–0 suture. MTJ transection results in a transient decrease in loading, with reintegration of the MTJ observed by D7-10 post-surgery. Following sterilization of the surgery region, a small incision was made on the posterior surface of the hind paw, the FDL tendon was located and completely transected using micro spring-scissors. The tendon was repaired using an 8-0 suture and the skin was closed with a 5-0 suture. Following surgery, animals resumed prior cage activity, food intake, and water consumption.

### Paraffin histology and immunofluorescence

Hind paws from *Scx-Cre*^*ERT2,Ai9*^*;Scx*^*GFP*^ mice were harvested prior to tendon surgeries, as well as at D7, 14, 21, 28, 35, and 42 post-surgery. Additionally, hind paws from *Scx-Cre; DTR*^*F/+*^ were harvested at D28 post-surgery. All hind paws were fixed in 10% neutral buffered formalin (NBF) at room temperature for 72 hr and were subsequently decalcified in Webb Jee EDTA (pH 7.2–7.4) for 14 days at room temperature, processed, and embedded in paraffin. Five-micron sagittal sections were utilized for analysis. For immunofluorescence staining, *Scx-Cre*^*ERT2,Ai9*^*;Scx*^*GFP*^ sections were stained with GFP (1:500, Cat#: MA5-15256, INVITROGEN, Waltham, MA), tdTomato (1:500, Cat#: AB8181, SICGEN, Cantanhede, Portugal), and α-SMA-FITC (1:500, Cat#: F3777, Sigma Life Sciences, St. Louis, MO). *Scx-Cre; DTR*^*F/+*^ repair sections were stained with Postn (1:300, Cat#: <u>AB21519</u>9, ABCAM, Cambridge, United Kingdom). All sections were counterstained with the nuclear DAPI stain and imaged with a VS120 Virtual Slide Microscope (Olympus, Waltham, MA). Additionally, *Scx-Cre; DTR*^*F/+*^ repair sections were stained with Alcian blue/hematoxylin and Orange G (ABHOG) to assess tissue morphology and collagen deposition. Finally, *Scx-Cre; DTR*^*F/+*^ repair sections were further imaged using Second Harmonic Generation (SHG) imaging to facilitate collagen organization and deposition.

### Quantification of fluorescence

Fluorescent images scanned by the virtual slide scanner were quantified using Visiopharm image analysis software v.6.7.0.2590 (Visiopharm, Hørsholm, Denmark). Automatic segmentation via a threshold classifier was used to define and quantify specific cell populations based on cell number. An ROI was drawn to encapsulate both the scar tissue and tendon stubs. The number of fluorescent cells was quantified and normalized to the number of the total cell number (DAPI) to determine percentages of each cell type. An n = 3-5 was used for quantification of *Scx*^*Lin*^ and *Scx*^*GFP*^ cells. An n=5-7 was used for quantification of αSMA^+^ cells.

### Quantification of biomechanical properties

FDL tendons at D28 post-surgery from DTR and WT groups were harvested from the hind paws. Under a dissecting microscope, each tendon was carefully separated at the myotendinous junction (MTJ). The tarsal tunnel was cut and the tendon was released from the tarsal tunnel, isolated until the bifurcation of the digits and then cut and released. Any additional connective tissues (e.g., muscle) were removed under a dissecting microscope, and the tendon was prepared for uniaxial testing. Sandpaper was placed on each end of the tendon and glued together using cyanoacrylate (Superglue, LOCTITE). The tendon was periodically submerged in PBS to avoid any potential tissue drying. Next, gripped tendons were transferred into a semi-customized uniaxial microtester (eXpert 4000 MicroTester, ADMET, Inc., Norwood MA) [21–24]. The microtester was transferred to an inverted microscope (Olympus BX51, Olympus) to visualize the tendon and quantify the gauge length, width, and thickness. The gauge length of each sample was set as the end-to-end distance between opposing sandpaper edges and was set the same for all samples tested. The cross-section of the tendon was assumed as an ellipse, where the width and thickness of the tissue represents the major axis and the minor axes, respectively. Based on the measured width and thickness of each tendon, the area of the elliptical cross-section was computed. A uniaxial displacement-controlled stretching of 1% strain per second until failure was applied. Load and grip-grip displacement data were recorded and converted to stress-strain data, and the failure mode was tracked for each mechanically tested sample. Stiffness and peak load were calculated based on load-displacement curves, while tangent modulus and peak stress were calculated based on stress-strain curves. Note that this method of computing tangent modulus assumes that stress and strain are uniform within each specimen.

### RNA extraction, next generation sequencing (NGS), and data analysis of RNA-seq

FDL tendons (1 tendon per biological sample; N=3 biological samples per genotype) were harvested at D28 post-surgery and flash frozen in liquid nitrogen. Total RNA was isolated using the Powermasher II (DIAGNOCINE) to homogenize the tissue. RNA was isolated from the extract using Trizol (Life Technologies, Carlsbad, CA) and the RNeasy Plus Micro Kit (Qiagen, Valencia, CA) per manufacturer’s recommendations. The total RNA concentration was determined with the NanoDrop 1000 spectrophotometer (NanoDrop, Wilmington, DE) and RNA quality assessed with the Agilent Bioanalyzer (Agilent, Santa Clara, CA). The RNA integrity number (RIN) for all harvested samples was 7.73 ± 0.22 (mean ± standard deviation). The TruSeq Stranded mRNA Sample Preparation Kit (Illumina, San Diego, CA) was used for next-generation sequencing library construction per manufacturer’s protocols. Briefly, mRNA was purified from 200 ng total RNA with oligo-dT magnetic beads and fragmented. First-strand cDNA synthesis was performed with random hexamer priming followed by second-strand cDNA synthesis using dUTP incorporation for strand marking. End repair and 3’ adenylation was then performed on the double-stranded cDNA. Illumina adaptors were ligated to both ends of the cDNA and amplified with PCR primers specific to the adaptor sequences to generate cDNA amplicons of approximately 200–500 bp in size. The amplified libraries were hybridized to the Illumina flow cell and single end reads were generated for each sample using Illumina NovaSeq6000. The generated reads were demultiplexed using bcl2fastq version 2.19.0. Data cleaning and quality control was accomplished using FastP version 0.20.0. Read quantification was accomplished using subread-1.6.4 package (featureCounts). Data normalization and differential expression analysis of DTR relative to WT at a given time point was performed using DESeq2-1.22.1 with an adjusted p-value threshold of 0.05 on each set of raw expression measures. The ‘lfcShrink’ method was applied, which moderates log2 fold-changes for lowly expressed genes. DeSeq2 data was uploaded to *Enrichr* (https://maayanlab.cloud/Enrichr/) to perform pathway analysis of the DEGs in DTR vs WT. Additionally, StringDB (https://string-db.org/) was utilized to identify potential interactions between the DEGs.

### Statistical analysis

Experimental N was determined based on previously published work [7–9]. GraphPad Prism was utilized to analyze quantitative data and is presented as mean ± standard deviation (Stdev). Student’s t-test was utilized to compare between DTR vs WT groups. Mice were randomly selected for specific experimental outcome metrics prior to surgery and quantitative data (ex. fluorescence quantification, biomechanics) were analyzed in a blinded manner. p values ≤ 0.05 were considered significant. * indicates p<0.05, ** indicates p<0.01, *** indicates p<0.001, **** indicates p<0.0001.

## Results

### *Scx*^*Lin*^ cells continuously expand during the proliferative phase of healing tendons with additonal cells expressing *Scx*

To understand the dynamics and contributions of *Scx*^*Lin*^ cells during both tendon homeostasis and healing, we utilized the inducible *Scx-Cre*^*ERT2,Ai9*^; *Scx-GFP* **(Fig. 1A)** mouse model. During homeostasis, *Scx*^*LinAdult+*^*;Scx*^*GFP+*^ cells were the predominant population (65.3% of total cells), followed by *Scx*^*LinAdult-*^*;Scx*^*GFP-*^*cells* (27.51% of total cells), *Scx*^*LinAdult-*^*;Scx*^*GFP+*^ cells (4.72% of total cells), and finally *Scx*^*LinAdult+*^*;Scx*^*GFP-*^ cells (2.47% of total cells) **(Fig. 1B, C; Suppl. Fig. 1B, table 1)**. These data suggests that indeed the cell composition of homeostatic FDL tendons is heterogeneous with *Scx*^*LinAdult+*^*;Scx*^*GFP+*^ cells being the predominant population

**Figure 1.**
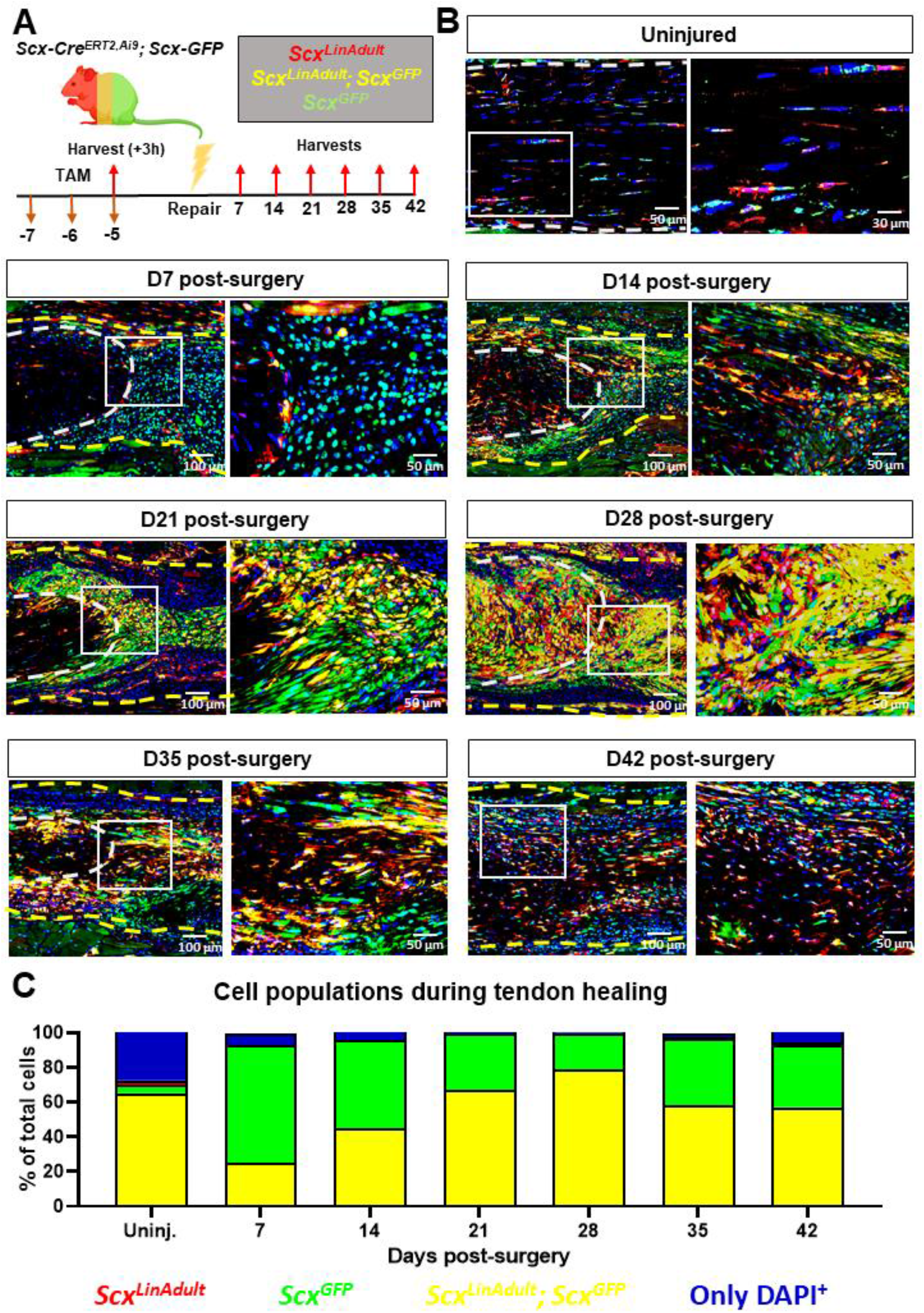
*Scx*^*Lin*^ cells continuously expand with additonal cells expressing *Scx* during tendon healing. **A**. Schematic of the mouse model used and timeline for tamoxifen injections, tendon surgeries, and tissue harvesting. **B**. Hind paws from *Scx-Cre*^*ERT2, Ai9*^; *Scx-GFP* were probed Red Fluorescence Protein (RFP), Green Fluorescence Protein (GFP), and were counterstained with the nuclear dye DAPI. **C**. Average cell density of *Scx*^*Lin+*^, *Scx-GFP*^*+*^, *Scx*^*Lin+*^*;Scx-GFP*^*+*^ and *Scx*^*Lin-*^*;Scx-GFP*^*-*^ (*DAPI*^*+*^). White dashed lines represent the tendon stub. Yellow dashed lines represent the bridging tissue between the tendon stubs. per timepoint. N=3-5 per timepoint. Significance was set to p<0.05.

By D7 post-surgery, a substantial shift in the composition of the tenocyte cell enviornment was observed, with *Scx*^*LinAdult-*^*;Scx*^*GFP+*^ cells being the predominant population (67.4% of total cells), while *Scx*^*LinAdult+*^*;Scx*^*GFP+*^ cells accounted for 25.24% of the total cells **(Fig. 1B, C; Suppl. Fig. 1B, table 1)**. Interestingly, there was a continuous decrease in the proportion of *Scx*^*LinAdult-*^*;Scx*^*GFP+*^ cells from D7 (67.4% of total cells) to D28 (21.03% of total cells) post-surgery. In parallel, the *Scx*^*LinAdult+*^*;Scx*^*GFP+*^ population continuously expanded from D7 (25.24% of total cells) to D28 (78.88% of total cells) post-surgery. Finally, the overall *Scx*^*Lin*^ pool (defined as the sum of *Scx*^*LinAdult+*^*;Scx*^*GFP+*^ and *Scx*^*LinAdult+*^*;Scx*^*GFP-*^) was substantially increased from homeostasis (72.48% of total cells) in response to injury at D28 post-surgery (99.92% of total cells) **(Fig. 1B, C; Suppl. Fig. 1B, table 1)**.

From D35 through D42 post-surgery, the proportion of cells that were *Scx*^*LinAdult+*^*;Scx*^*GFP+*^ progressively decreased (D35:58.49%; D42: 57.18% of total cells) toward the levels observed during homeostasis (65.23% of total cells) **(Fig. 1B, C; Suppl. Fig. 1B, table 1)**. In addition, at D35 and D42 there was an expansion of the *Scx*^*LinAdult-*^*;Scx*^*GFP-*^ population **(Fig. 1B, C; Suppl. Fig. 1B, table 1)**. Taken together, these data suggest that during the late stages of healing, the cell composition of the healing tendon begins to shift back toward the composition that is observed during homeostasis.

### *Scx* expression is required for maintenance of myofibroblast (*αSMA*^+^) cells during tendon healing

We have previously shown that *αSMA*^*+*^ myofibroblasts are primarily derived from the broad *Scx*^*Lin*^ pool during adult tendon healing [9]. To get a better understanding of the dynamics of *αSMA*^*+*^ myofibroblast presence and their relationship to active *Scx* expression and the adult *Scx*^*Lin*^ pool during healing, we utilized the inducible *Scx-Cre*^*ERT2,Ai9*^; *Scx-GFP* **(Fig. 2A)** mouse model and tracked the dynamics between *aSMA*^*+*^ and *Scx*^*GFP+*^ **(Fig. 2B)** as well as *aSMA*^*+*^ and adult *Scx*^*Lin+*^ cells between D7 until D42 post-surgery **(Suppl. Fig. 2A, B)**.

**Figure 2.**
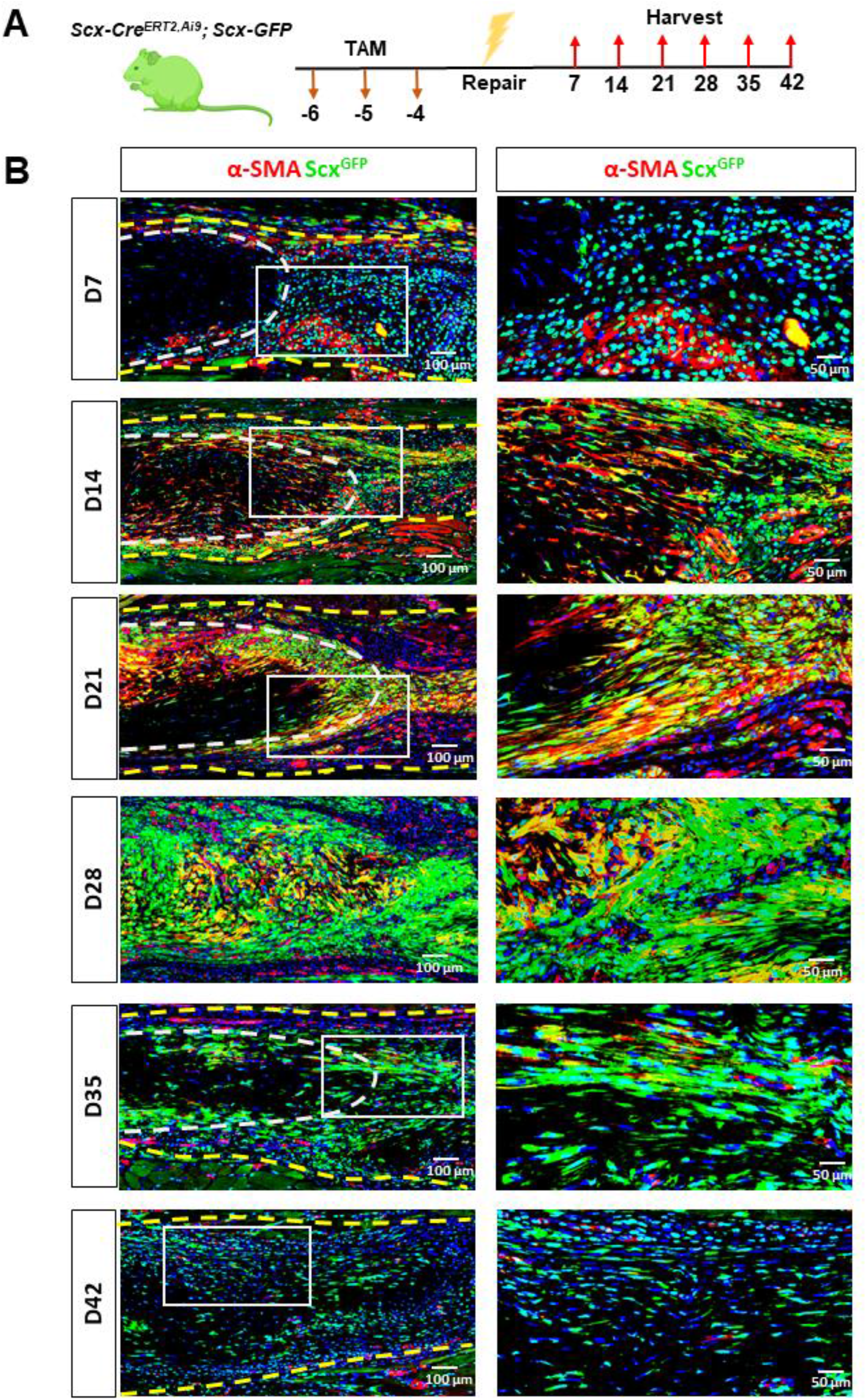
A-SMA myofibroblasts are transiently expressed between D14 and D35 post-surgery and retain *Scx*^*+*^ expression. **A**. Schematic of the mouse model used and timeline for tamoxifen injections, tendon surgeries, and tissue harvesting. **B**. Hind paws from the *Scx-Cre*^*ERT2,Ai9*^; *Scx-GFP* mice were probed Red Fluorescence Protein (RFP), Green Fluorescence Protein (GFP), and were counterstained with the nuclear dye DAPI.

By D7 post-surgery, there were virtually no *αSMA*^*+*^ myofibroblasts in the healing tendons **(Fig. 2B)**. However, from D14 to D21, there was a substantial increase in *αSMA*^*+*^ cells in the healing tendon with nearly all *αSMA*^*+*^cells actively expressing *Scx* **(Fig. 2B)**. At D28 and D35 post-surgery, *aSMA*^*+*^ cells persisted, and *Scx* expression was retained, however, the presence of *aSMA*^*+*^ was markedly reduced relative to D14 and D21 **(Fig. 2B)**. Finally, by D42, there were no *aSMA*^*+*^ cells in the injured tendons **(Fig. 2B)**. In terms of the adult *Scx*^*Lin*^ contribution to myofibroblast fate, some *aSMA*^*+*^cells were derived from the adult *Scx*^*Lin*^ population between D14 and D35 post-surgery **(Suppl. Fig. 2A, B)**, though this population represents only a fraction of myofibroblasts.

### Healing DTR tendons exhibit deficits in structural mechanical and material properties

To define the requirement of the broad *Scx*^*Lin*^ cell pool during the proliferative phase of tendon healing, we utilized the *Scx-Cre*^*+*^; *Rosa-DTR*^*F/+*^ (DTR) mice to deplete *Scx*^*Lin*^ cells between D14 and D18 post-surgery. For controls, we utilized the *Scx-Cre*^*-*^; *Rosa-DTR*^*F/+*^ (WT). All mice received 5 consecutive local injections of Diphtheria Toxin (DT) in the injured tendon area and were harvested at D28 post-surgery **(Fig. 3A)**.

**Figure 3.**
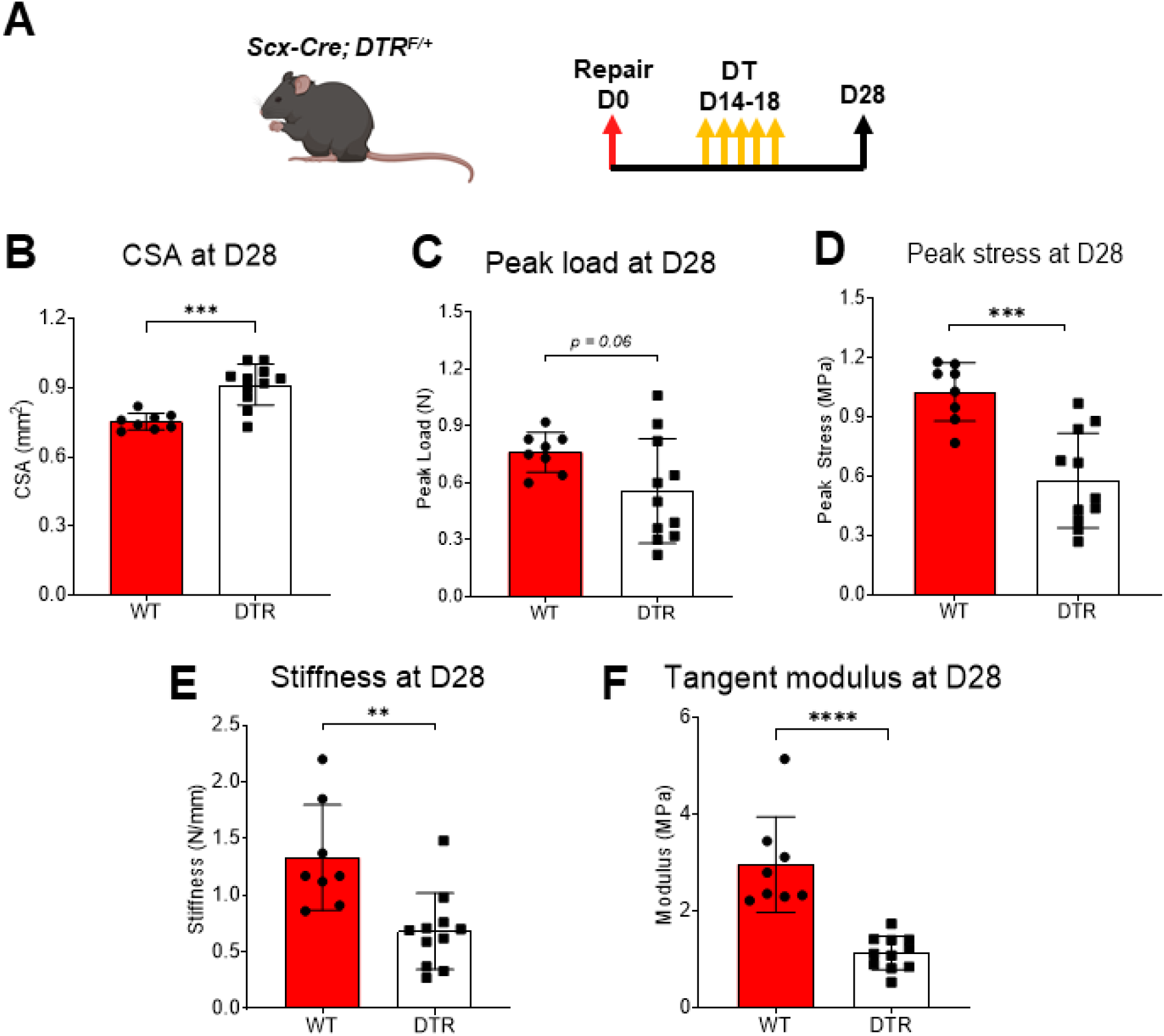
DTR tendons exhibit significant deficits in biomechanical properties. **A**. Schematic of the mouse model used and timeline for tendon surgeries, DT injections, and tissue harvesting. CSA **(B)**, peak load **(C)**, peak stress **(D)**, stiffness **(E)**, and tangent modulus **(F)** of the the D28 DTR vs WT tendons. Student’s t-test was utilized for statistical testing. N=8-12 per genotype. **:p<0.01; ***:p<0.001; ****:p<0.0001.

To assess whether *Scx*^*Lin*^ cells are required to restore tendon biomechanical properties, we quantified structural mechanical properties (peak load, stiffness) and material properties (peak stress, tangent modulus) **(Fig. 3B-F)**. DTR tendons exhibited a 21.33% (p<0.001) increase in CSA compared to WT littermates **(Fig. 3B)**. The peak load of DTR tendons exhibited a trending decrease (p=0.06) compared to WT **(Fig. 3C)**. The peak stress of DTR tendons was significantly decreased by 43.63% (p<0.001) compared to WT littermates **(Fig. 3D)**. As for stiffness, DTR tendons demonstrated a significant 48.72% decrease (p<0.01) compared to WT **(Fig. 3E)**. Finally, DTR tendons had a 61.94% (p<0.0001) decrease in tangent modulus compared to WT repairs **(Fig. 3F)**. Collectively, these data suggest that *Scx*^*Lin*^ cells are required for restoration of tendon biomechanics during healing.

### Bulk RNA-seq reveals enrichment of pathways related to ECM synthesis and organization, cell-matrix adhesion, and cellular proliferation in DTR tendons

To better define the biological mechanisms associated with impaired healing in DTR tendon repairs, we performed bulk RNA-seq and assessed changes in the transcriptomic profile between DTR and WT tendons at D28. *Scx*^*Lin*^ cell depletion between D14-18 post-surgery resulted in a significant transcriptomic shift of healing tendons with 197 genes significantly increased and 269 genes significantly decreased in DTR compared to WT tendons **(Fig. 4A-C)**.

**Figure 4.**
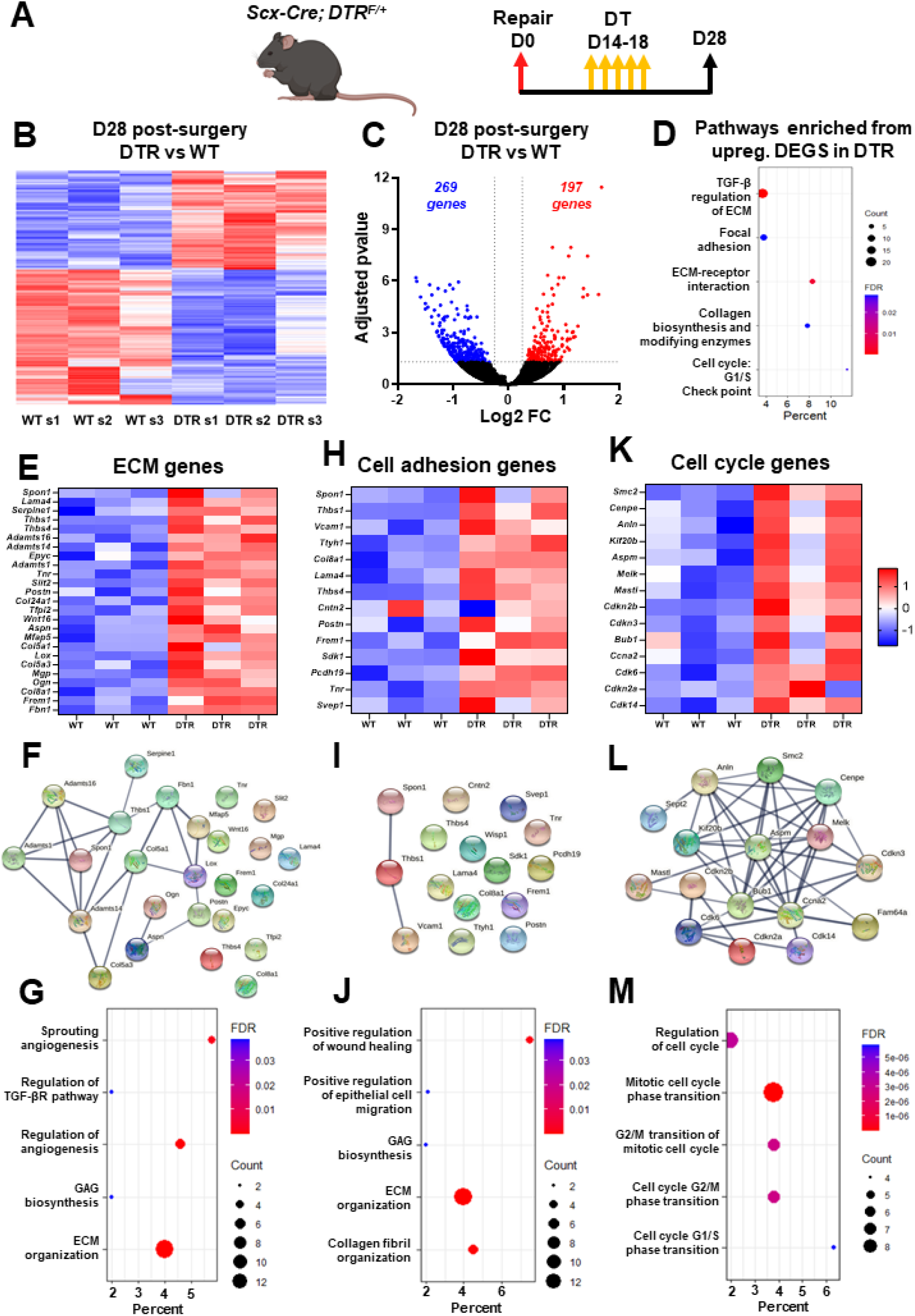
DTR tendons exhibit enriched biological pathways related to ECM synthesis and organization, cell-ECM receptor interaction, and cellular mitosis/proliferation. **A**. Schematic of the mouse model used and timeline for tendon surgeries, DT injections, and tissue harvesting. **B**. Heatmap of all significantly different genes between D28 DTR and WT tendons. **C**. Volcano plot of all the significantly different genes between D28 DTR and WT tendons. **D**. Enriched biological pathways from the upregulated genes in D28 DTR relative to WT tendons. **E**. Heatmap with all the ECM **(E)**, cell adhesion **(F)**, and cell cycle **(G)** genes significantly upregulated in D28 DTR tendons. **H**. Protein-protein communication of all the ECM **(H)**, cell adhesion **(I)**, and cell cycle **(J)** genes significantly upregulated in D28 DTR tendons.

GO analysis via *Enrichr* identified multiple biological pathways that were significantly enriched in the DTR tendons based on the upregulated genes **(Fig. 4D)**. In specific, there were three main pathways enriched related to ECM (*TGF-β regulation of ECM* and *Collagen biosynthesis and modifying enzymes)*, cell-ECM adhesion (*Focal adhesion, ECM-receptor interaction)*, and cell mitosis/proliferation (*Cell cycle: G1/S Check point)* **(Fig. 4D)**.

Genes related to each enriched pathway were identified after screening all the significantly upregulated genes in DTR tendons using *StringDB*. For the ECM-related pathway, a total of 25 genes were identified and were expressed at significantly higher levels in all three different biological groups in DTR compared to WT tendons **(Fig. 4E)**. To assess any potential interactions between the ECM-related genes, *StringDB* was utilized and it was found that many of those genes do interact with each other (e.g. *Serpine1, Adamts16, Fbn1, Mfap5, Lox, Col5a1, Thbs1, Adamts1, Spon1, Adamts14, Col5a3, Postn, Aspn*, and *Ogn)* while other ECM-related genes did not exhibit interactions with the rest of the genes (e.g. *Thbs4, Epyc, Col8a1, Col24a1, Lama4*) **(Fig. 4F)**. Finally, GO analysis identified pathways related to *angiogenesis, regulation of TGFβ receptor pathway, biosynthesis of glycosaminoglycans (GAGs)*, and *ECM organization* were significantly enriched in DTR repairs **(Fig. 4G)**. Regarding the second main pathway enriched (*cell-ECM adhesion*, ***Fig. 4D)***, a total of 14 genes were identified **(Fig. 4H)**. Analysis of the interaction of those proteins with each other showed that only *Spon1, Thbs1*, and *Vcam1* seem to have some interaction with each other **(Fig. 4I)**. GO analysis identified pathways related to *positive regulation of wound healing and epithelial cell migration, GAG biosynthesis, ECM organization*, and *collagen fibril organization* significantly enriched in DTR tendons **(Fig. 4J)**. Finally, for *cell mitosis/proliferation* (**Fig. 4D**), screening using *StringDB* identified a total of 14 genes **(Fig. 4K)**. Interaction analysis demonstrated that all the cell mitosis related genes interact with each other, suggesting that they all follow the same mechanisms for cellular proliferation **(Fig. 4L)**. GO analysis identified pathways related to *regulation of cell cycle, mitotic cell cycle phase transition, G1/S and G2/M transition of mitotic cell cycle* were significantly enriched for DTR tendons **(Fig. 4M)**.

Taken together, these data suggest that D28 DTR tendons actively synthesizing ECM molecules related to wound closure. In parallel, there is cell proliferation (potentially of *Scx*^*Lin*^ cells [25]) in the healing DTR tendons that may be an attempt to increase cell density and to further aid in the wound closure. Such pathways are usually enriched when a tissue is within the proliferative healing phase [26–29]. Considering that *Scx*^*Lin*^ cell depletion was performed between D14-18 post-surgery, which is firmly in the proliferative phase, these data suggest that there is a stagnation of the healing response around the time of depletion (D14-18).

To validate that the healing response of D28 DTR tendons was stalled in a manner consistent with the molecular program of the proliferative phase, we conducted additional analysis of bulk RNA-seq data from our previously published study [7] from WT tendons at D14 post-surgery (proliferative phase) compared to WT at D28 post-surgery (remodeling phase) **(Suppl. Figure 2)**. GO analysis of enriched pathways and biological processes based on all genes that were significantly increased in the D14 post-surgery (proliferative phase), showed that pathways and biological processes related to ECM synthesis and organization, cell-ECM adhesion, and cell proliferation/ mitosis were significantly activated **(Suppl. Figure 2A-J)**.

### Bridging matrix of DTR tendons exhibits less mature collagen fibrils and high amount of proteoglycans and glycoproteins

To better understand the morphological features that occur concomitant with mechanical deficits and transcriptional shifts in DTR healing tendons, we performed Alcian Blue Hematoxylin Orange Green (ABHOG) staining and it revealed that indeed, the bridging tissue of D28 DTR tendons exhibited an impaired matrix quality compared to D28 WT tendons **(Fig. 5B)**. More specifically, DTR tendons exhibited thinner and more immature collagen fibrils, and increased staining of proteoglycans/glycoproteins (PGs/GPs) compared to WT tendons **(Fig. 5B)**, consistent with our bulk RNA-seq data **(Fig. 4)**. SHG imaging further validated the absence of mature collagen fibrils in DTR tendons relative to WT repairs **(Fig. 5C)**. Periostin (Postn) was among the significantly upregulated PGs/GPs in DTR tendons repairs **(Fig. 4E, H)**. Immunofluorescence demonstrated that increased Postn expression occurred in DTR tendon repairs relative to WT **(Fig. 5D)**. Taken together, these data suggest that *Scx*^*Lin*^ cells are required during the proliferative healing phase for matrix synthesis and maturation/remodeling, and their absence substantially impairs the composition, structure, and function of during tendon healing.

**Figure 5.**
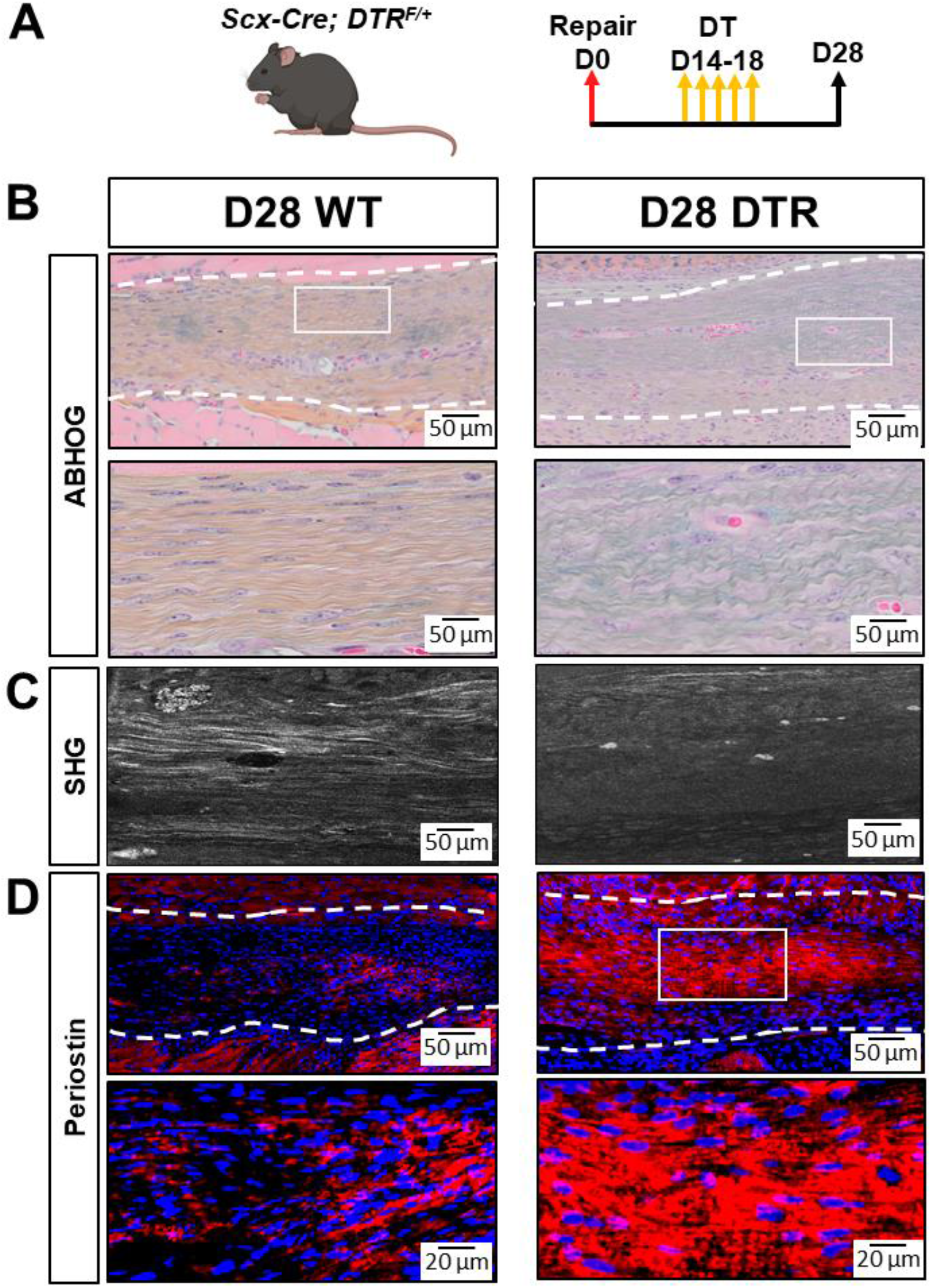
DTR tendons exhibit an immature bridging matrix tissue at D28 post-surgery. **A**. Schematic of the mouse model used and timeline for tendon surgeries, DT injections, and tissue harvesting. **B**. ABHOG staining to visualize the structure and organization of the healing DTR vs WT tendons. **C**. SHG imaging to visualize mature collagen fibrils in the healing DTR vs WT tendons. **D**. Periostin staining of D28 DTR vs WT tendons. N=3 per genotype.

### DTR tendons fully restore their biomechanical properties by D56 post-surgery

To investigate whether the structure-function deficits during healing following *Scx*^*Lin*^ depletion (at D14-18) are transient or permanent, healing tendons were harvested for biomechanical testing at D56 post-surgery (late remodeling healing phase) **(Fig. 6A)**.

**Figure 6.**
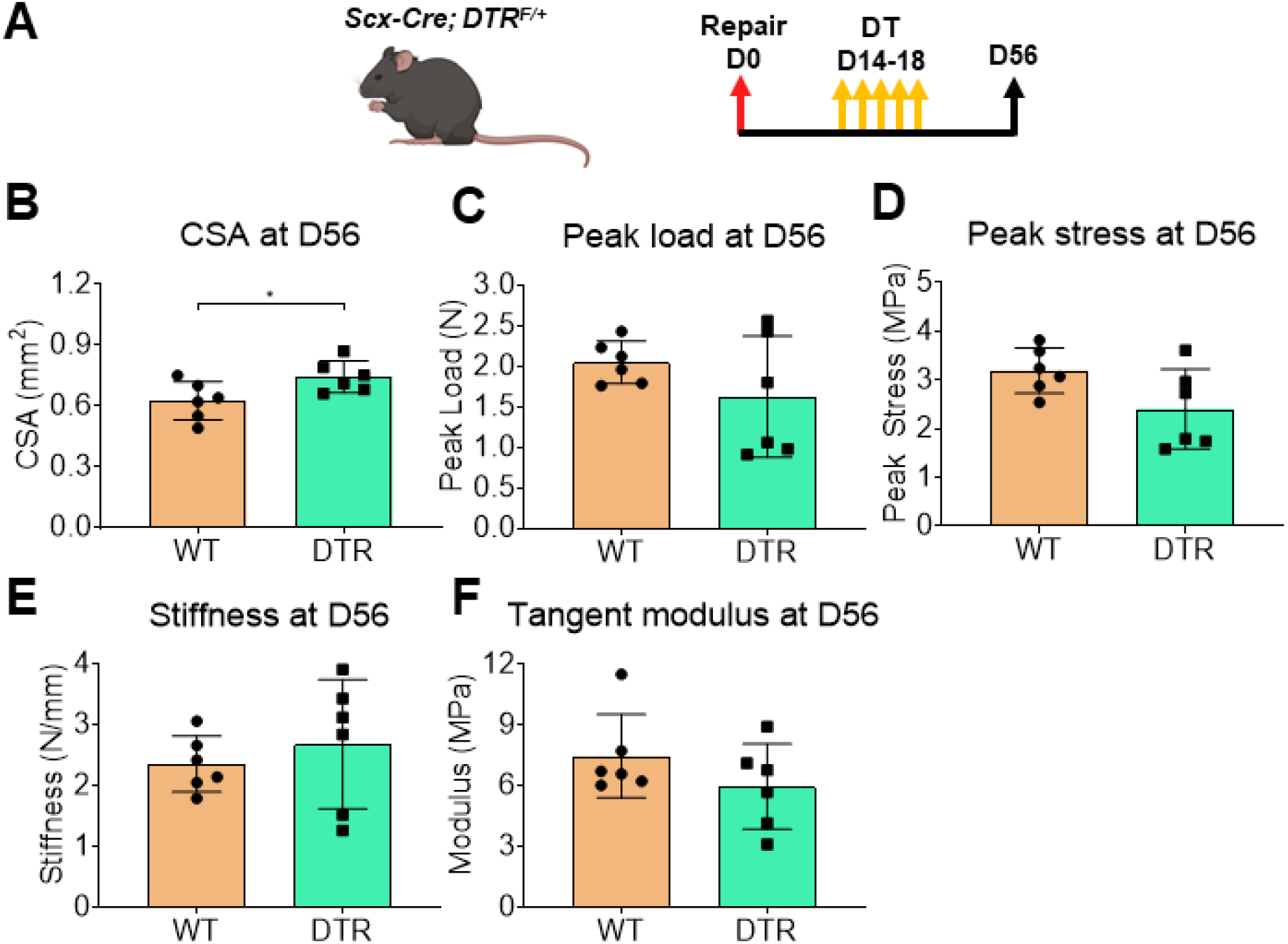
DTR tendons fully restore biomechanical propertes by D56 post-surgery. **A**. Schematic of the mouse model used and timeline for tendon surgeries, DT injections, and tissue harvesting. CSA **(B)**, peak load **(C)**, peak stress **(D)**, stiffness **(E)**, and tangent modulus **(F)** of the the D56 DTR vs WT tendons. Student’s t-test was utilized for statistical testing. N=6 per genotype. **:p<0.01; ***:p<0.001; ****:p<0.0001.

DTR tendons exhibited a 17.74% (p<0.05) increase in CSA compared to WT littermates **(Fig. 6B)**. The peak load, peak stress, stiffness, and tangent modulus of DTR tendons were not significantly different (p>0.05) compared to WT **(Fig. 6C-F)**. These data suggest that in the long-term DTR tendons are able to restore structural mechanical and material properties during healing, however the retained increase in CSA suggests a potential lack in further remodeling of the scar tissue.

## Discussion

A comprehensive understanding of the different cell populations and their associated molecular mechanisms during tendon healing is pivotal to design targeted therapeutics for tendon regeneration. Here, we first characterized the dynamics of two major subpopulations of the broad *Scx*^*Lin*^ cell pool (adult *Scx*^*Lin*^ and *Scx*^*GFP*^ cells) during tendon healing, and established that new *Scx*^*+*^ cells are continuously added to the overall *Scx*^*Lin*^ pool from the early proliferative (D7) to the remodeling phase (D28) of healing. Next, we also demonstrated the dynamics αSMA^+^ myofibroblasts and their relationship to *Scx*^*GFP*^ cells throughout healing. Strikingly, almost all αSMA myofibroblasts retained *Scx* expression for the duration of myofibroblast presence during healing, suggesting a potential requirement for Scx expression to maintain myofibroblast fate. Next, we investigated the function of a subset of the broad *Scx*^*Lin*^ pool by inducibly depleting *Scx*^*Lin*^ cells during the proliferative phase and found that *Scx*^*Lin*^ cells are required for ECM synthesis, organization, and restoration of tendon biomechanics. Strikingly, we found that *Scx*^*Lin*^ depletion from D14-18 caused a temporary stagnation of the healing response, stalling DTR tendons in the proliferative phase of healing even at D28 post-surgery. However, this stagnation was temporary such that by D56, mechanical properties were restored to the same level as WT repairs, suggesting that DTR tendons underwent successfully through the remodeling phase.

Disrupting tendon homeostasis via acute injury and repair resulted in significant shifts in the cell environment. By D7, *Scx*^*LinAdult-*^; *Scx*^*GFP+*^ cells that expressed *Scx* in response to injury, were the predominant population in the healing tendons. Although their source is not clear, it could be from the *Scx*^*LinAdult-*^*;Scx*^*GFP-*^ population that is present during homeostasis. Indeed, recent findings from our lab and others have identified multiple tenocytes during homeostasis that do not express *Scx* but express markers related to wound healing and ECM synthesis [13, 14, 25, 30]. Another alternative source could be epitenon-derived cells that migrate into the healing tendon. Indeed, we previously identified multiple *Scx*^*-*^ epitenon cell subpopulations in healthy FDL tendons [13].

*Scx*^*LinAdult-*^; *Scx*^*GFP+*^ cells were identified as the predominant population between D7 and D14. Previously, we have shown that during these timepoints, the initital bridging tissue that connects the tendon stubs is formed by *Scx*^*Lin*^ cells [8, 25]. From D14 to D28 post-surgery (proliferative and remodeling phase), the proportion of *Scx*^*LinAdult-*^; *Scx*^*GFP+*^ cells was reduced, while the *Scx*^*LinAdult+*^; *Scx*^*GFP+*^ population continuously expanded. Taken together, these data suggest that *Scx*^*LinAdult-*^; *Scx*^*GFP+*^ and *Scx*^*LinAdult+*^; *Scx*^*GFP+*^ cells may have distinct functions during tendon healing. More specifically, *Scx*^*LinAdult-*^; *Scx*^*GFP+*^ cells may be responsible for synthesizing the initial bridging scar tissue. In contrast, *Scx*^*LinAdult+*^; *Scx*^*GFP+*^ cells may be responsible for remodeling and organization of the bridging scar tissue.

Myofibroblasts demonstrated a transient presence, peaking in the proliferative phase (D21), decreasing during the remodeling phase (D28 through D35), and fully resolving by the later remodeling phase (D42). During their presence in healing tendons, almost all myofibroblasts were *Scx*^*+*^ and retained their *Scx* expression until they were resolved. This intriguing finding suggests that *Scx* expression is required for αSMA myofibroblast maintenance during tendon healing. In support to this finding, previous studies in cardiac fibroblasts have shown that *Scx* expression is both required and necessary for the synthesis of αSMA protein and conversion of fibroblasts to myofibroblasts [31–34]. In specific, Bagchi *et al*., utilized an *in vitro* system for *Scx* gain- or loss- of function in primary cardiac fibroblasts and their effect on αSMA expression [34]. They found that *Scx* knockdown reduced αSMA expression and incorporation into stress fibers while overexpression of *Scx* resulted in further induction of αSMA expression and production of stress fibers [34]. Given the central role of *Scx* expression for αSMA presence and maintenance, the transcription factor *Scx* could be a potential therapeutic target for fibrosis.

To understand the requirements for the broad *Scx*^*Lin*^ cell population during the proliferative healing phase, we depleted these cells between D14 and D18 post-surgery and found that these cells are important for restoration of structure, composition, and function in healing tendons. In specific, the bridging matrix tissue of D28 DTR tendons was characterized by immature collagen fibrils, increased expression of multiple ECM glycoproteins and proteoglycans, and reduced structural mechanical and material properties. These data suggest that *Scx*^*Lin*^ cells play a key role in both the synthesis and remodeling of the ECM in the bridging tissue. In accordance to our findings, multiple studies have previously assumed that *Scx*^*Lin*^ cells are required during tendon healing for ECM synthesis, organization, and wound clousure [8, 9, 16, 17, 35]. However, in this study, we directly tested and identified the requirements of *Scx*^*Lin*^ cells during tendon healing.

GO analysis from bulk RNA-seq at D28 post-surgery demonstrated that pathways and biological processes related to ECM synthesis and organization, cell-ECM adhesion, and cell proliferation/mitosis were significantly enriched in DTR tendons. Interestingly, such pathways and biological processess are typically upregulated when tissues are going through the proliferative healing phase [26–29]. Indeed, GO analysis of bulk RNA-seq data between WT D14 (proliferative phase) and WT D28 (remodeling phase) post-surgery timepoints, revealed that pathways and biological mechanisms related to ECM synthesis and organization, cell-ECM adhesion, and cell proliferation/mitosis were significantly enriched in the D14 post-surgery tendons. In addition to the transcriptomic data, both the morphology and biomechanics of the D28 DTR tendons were almost identical with those of D14 WT tendons that we have previously published in separate studies [7, 13, 21, 36]. Taking into account that *Scx*^*Lin*^ depletion was performed between D14-18 post-surgery, which is firmly in the proliferative phase, it suggests that with depletion, DTR tendons had become stagnant during healing, essentially stalling the healing process in the proliferative phase at least through D28 post-surgery. Next, we examined the long-term ability of DTR tendons to restore the healing process to that of WT. Indeed, at D56 post-surgery, DTR tendons were able to fully restore both their structural mechanical and material properties in the same levels as D56 WT tendons. This suggests that DTR tendons eventually progressed through the normal stages of healing including tissue remodeling, and suggests great plasticity of the healing process even when disrupted during the proliferative phase.

This study is not without limitations. First, although we distinguished between adult *Scx*^*Lin*^ and *Scx*^*GFP*^ cells during the different healing phases (D7 through D42 post-surgery), this depletion model targets both of these subpopulations. As such, it not possible to further decipher the requirements of specific subpopulations of cells in the overall *Scx*^*Lin*^ pool (e.g., roles of only the adult *Scx*^*Lin*^ cells) with this model. Previous efforts to deplete cells actively expressing *Scx* by using the inducible *Scx-Cre*^*ERT2*^ crossed to Rosa-DTA [37] mice resulted in insufficient recombination. Thus, in order to deplete specific subpopulations of *Scx*^*Lin*^ cells in a temporally-dependent manner, future work could utilize *Scx-Cre*^*ERT2*^;Rosa-DTR mice, although this model requires both TMX and DT administration, and discrepencies in labelling vs. depletion may be observed.

Taken together, in this study, we first demonstrated in detail the dynamics of two major subpopulations of the broad *Scx*^*Lin*^ pool (adult *Scx*^*Lin*^ and *Scx*^*GFP*^ cells) during the entire tendon healing course. We made the intriguing finding that *Scx* expression is required for myofibroblast maintenance and in parallel provided clear evidence of the spatial and temporal presence of active myofibroblasts during tendon wound healing. Next, we directly tested the role of *Scx*^*Lin*^ cells by depleting them in the proliferative healing phase and found that they are required to produce and remodel ECM, as well as to restore tendon biomechanics. Their absence resulted in a temporary stagnation of healing response, though, long-term, DTR tendons eventually healed to the same extent as WT repairs. Deciphering the time-dependent roles of *Scx*^*Lin*^ cells during tendon healing will better inform therapeutic target selection and promote regenerative tendon healing.

## Supplemental figures

**Supplemental figure 1.**
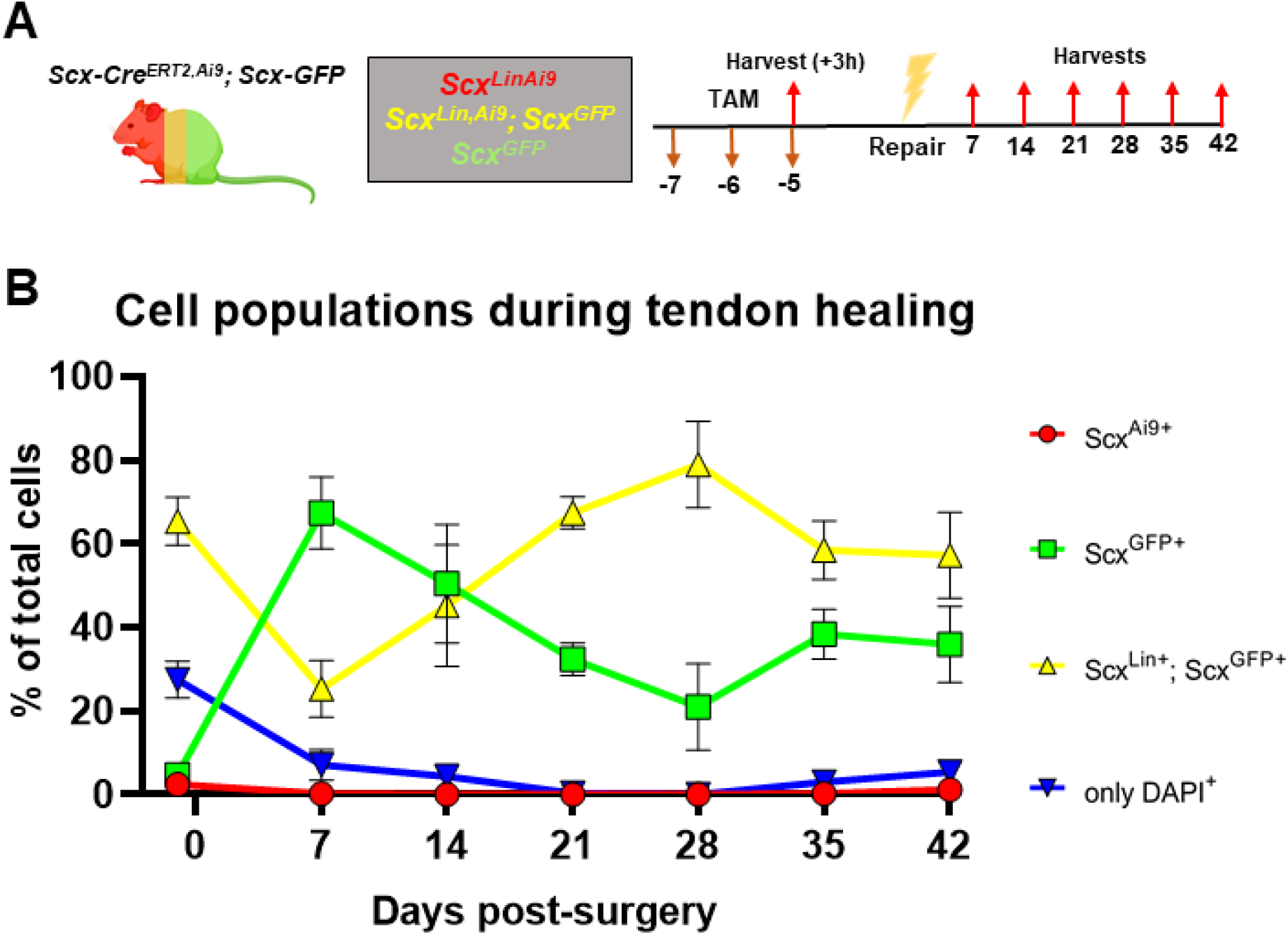
*Scx*^*Lin*^ cells continuously expand with additonal cells expressing *Scx* during tendon healing. **A**. Schematic of the mouse model used and timeline for tendon surgeries, DT injections, and tissue harvesting. **B**. Average cell density overtime of *Scx*^*Lin+*^, *Scx-GFP*^*+*^, *Scx*^*Lin+*^*;Scx-GFP*^*+*^ and *Scx*^*Lin-*^*;Scx-GFP*^*-*^ (*DAPI*^*+*^). N=3-5 per timepoint. One-way ANOVA with Tukey’s post-hoc was used to assess statistical significance of cell density between different cell populations (*Scx*^*Lin+*^, *Scx-GFP*^*+*^, *Scx*^*Lin+*^*;Scx-GFP*^*+*^ and *Scx*^*Lin-*^*;Scx-GFP*^*-*^). One-way ANOVA with Tukey’s post-hoc was used to assess statistical significance of cell density for each sub-populations at different timepoints (D-1 pre-surgery, D7, D14, D21, D28, D35, and D42 post-surgery).

**Supplemental figure 2.**
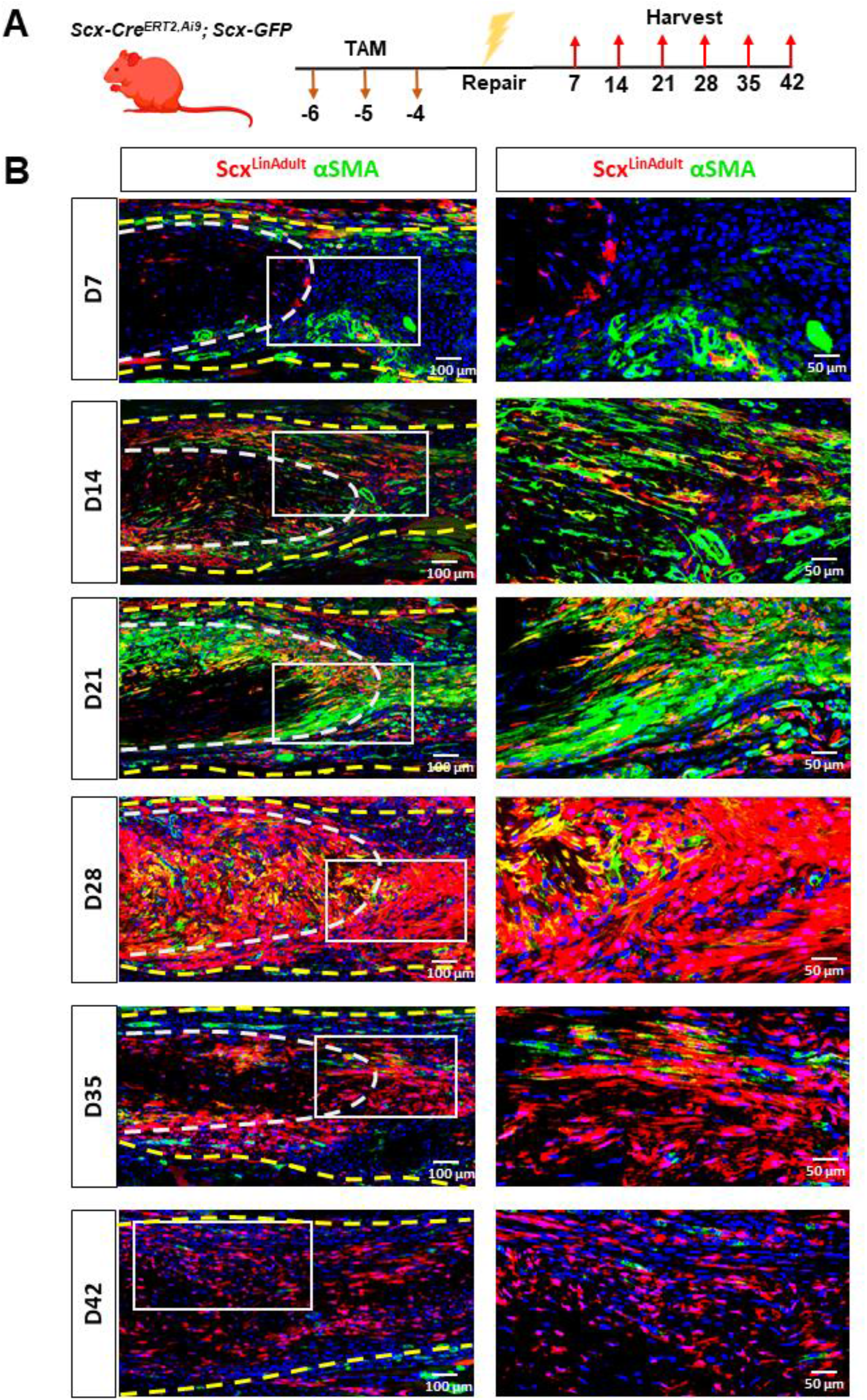
Dynamics between *Scx*^*Lin*^ cells and *α*SMA myofibroblasts. **A**. Schematic of the mouse model used and timeline for tamoxifen injections, tendon surgeries, and tissue harvesting. **B**. Hind paws from the *Scx-Cre*^*ERT2,Ai9*^; *Scx-GFP* mice were probed Red Fluorescence Protein (RFP) to visualize *Scx*^*Lin*^ cells and FITC to visualize αSMA myofibroblasts. All samples were counterstained with the nuclear dye DAPI. N=3-5 per timepoint.

**Supplement figure 3.**
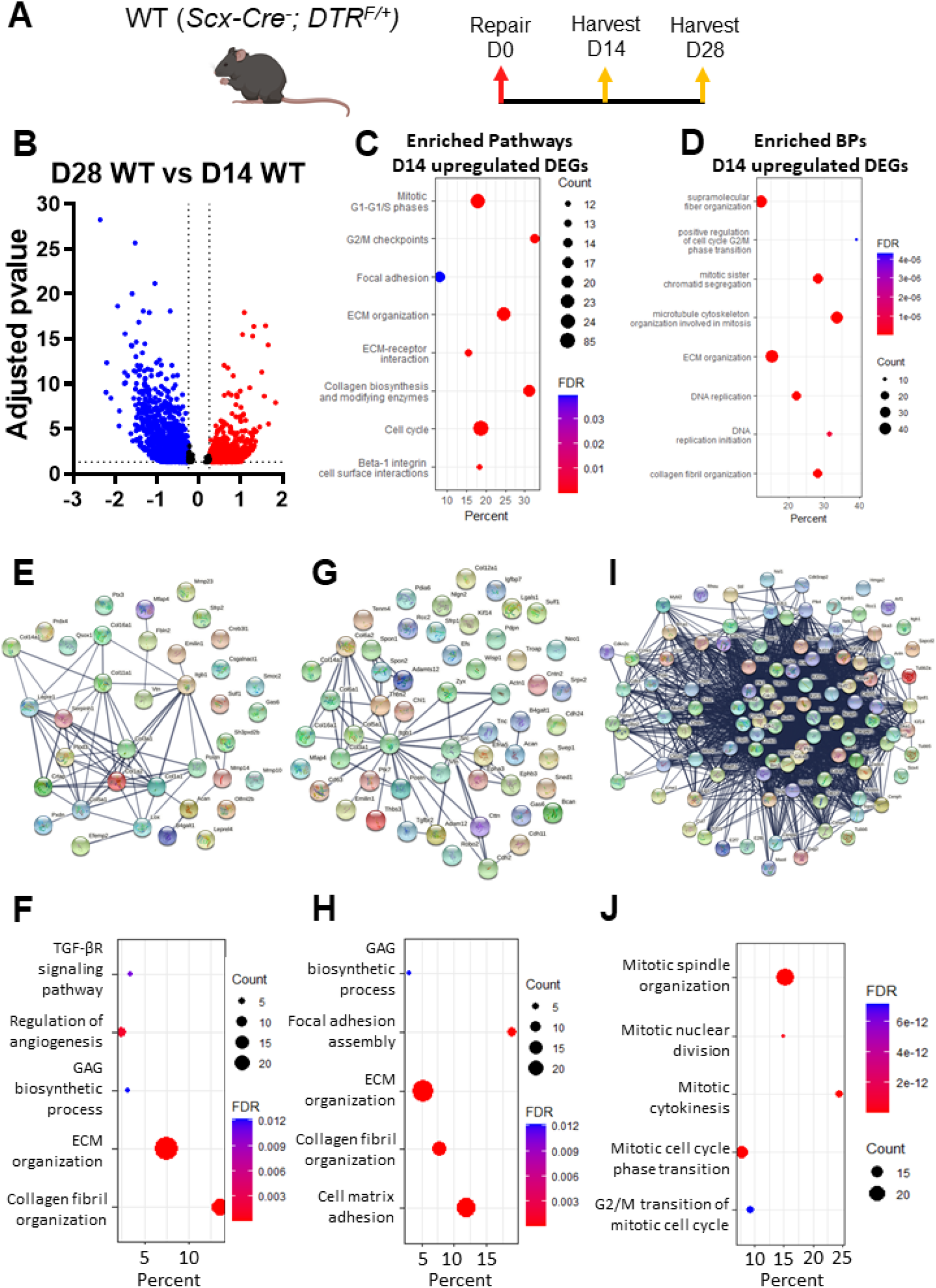
FDL tendons in the proliferative phase exhibit enriched biological pathways related to ECM synthesis and organization, cell-ECM receptor interaction, and cellular mitosis/proliferation compared to FDL tendons in the remodeling phase. **A**. Schematic of the mouse model used and timeline for tendon surgeries, DT injections, and tissue harvesting. **B**. Volcano plot of all the significantly different genes between D28 vs D14 WT tendons. Enriched pathways **(C)** and biological processess **(D)** between D28 vs D14 WT tendons. **E**. Protein-protein communication of all the ECM **(E)**, cell adhesion **(G)**, and cell cycle **(I)** genes between D28 vs D14 WT tendons. Enriched pathways related ECM **(F)**, cell adhesion **(H)**, and cell cycle **(J)** between D28 vs D14 WT tendons.

